# *Drosophila* PLP forms centriolar-clouds that promote centriole stability, cohesion and MT nucleation

**DOI:** 10.1101/161554

**Authors:** Helio Roque, Metta Pratt, Errin Johnson, Jordan W. Raff

## Abstract

Pericentrin is a conserved centrosomal protein whose dysfunction has been linked to several human diseases. The precise function of Pericentrin, however, is controversial. Here, we examine *Drosophila* Pericentrin-like-protein (PLP) function *in vivo*, in tissues that form both centrosomes and cilia. *PLP* mutant centrioles exhibit four major defects: (1) They are too short and have subtle structural defects; (2) They separate prematurely, and so overduplicate; (3) They organise fewer MTs during interphase; (4) They fail to establish and/or maintain a proper connection to the plasma membrane— although, surprisingly, mutant centrioles can still form an axoneme and recruit transition zone (TZ) proteins. We show that PLP helps to form “ pericentriolar clouds” of electron-dense material that emanate from the central cartwheel spokes and spread outward to surround the mother centriole. The partial loss of these structures may explain the complex centriole, centrosome and cilium defects we observe in *PLP* mutant cells.

## Introduction

Centrioles are complex MT-based structures that duplicate precisely once per cell cycle when a “ daughter” centriole is assembled on the side of a “ mother centriole during S-phase (Arquint and Nigg, 2016). In proliferating tissues, centrioles form centrosomes when they recruit pericentriolar material (PCM) around themselves (Conduit et al., 2015). The PCM contains several hundred proteins (Alves-Cruzeiro et al., 2013), including many involved in nucleating and organising MTs, so centrosomes function as the major MT organising centres (MTOCs) in many eukaryotic cells. In interphase, the centrioles organise relatively small amounts of PCM, but the PCM expands dramatically as cells enter mitosis in a process termed centrosome maturation (Palazzo et al., 2000; Conduit et al., 2015). In non-proliferating tissues, the centrioles often migrate to the cell cortex, where the mother centriole organises the assembly of a cilium. The cilium can be motile—so moving the cell, or generating liquid flow around the cell—or immotile, with mechano- and/or chemo-sensory functions. Defects in centriole, centrosome and cilium function have been linked to a wide range of human pathologies, including cancer, microcephaly, dwarfism and obesity (Nigg and Raff, 2009; Bettencourt-Dias et al., 2011

Pericentrin is one of the best studied centrosomal proteins, and it has been implicated in several aspects of centriole, centrosome and cilium function (Delaval and Doxsey, 2010). In vertebrates, Pericentrin interacts with several PCM proteins, and it appears to be an important driver of mitotic centrosome maturation (Zimmerman et al., 2004; Purohit et al., 1999; Haren et al., 2009;Lee and Rhee, 2011; Lawo et al., 2012). In flies, however, the Pericentrin-like-protein (PLP) has a more minor role in mitotic centrosome assembly (Martinez-Campos et al., 2004; Richens et al., 2015; Lerit et al., 2015), especially when compared to proteins like Cnn and Spd-2 (Cdk5Rap2/Cep215 and Cep192 in vertebrates, respectively) that form a scaffold upon which the mitotic PCM is assembled (Conduit et al., 2014; Feng et al., 2017). Instead, studies in fly cultured cells suggest that PLP helps to organise the interphase PCM (Mennella et al., 2012). PLP molecules are polarised around the mother centriole, with their C-termini linked to the centriole, and their N-termini extending outwards away from the centriole (Fu and Glover, 2012; Mennella et al., 2012). The MT nucleator γ-tubulin, and the mitotic scaffold proteins Cnn and Spd-2, are also recruited around the mother centriole during interphase, and this recruitment is abolished when PLP is depleted, although it has not been documented how this loss of interphase PCM influences the ability of these centrioles to organise MTs (Mennella et al., 2012). Interestingly, Pericentrin adopts a similarly polarised and extended conformation around the mother centriole in human cells (Lawo et al., 2012; Sonnen et al., 2012). Pericentrin also has several other functions in vertebrate cells: It is required for proper cilia function (Jurczyk et al., 2004); it is cleaved by Separase towards the end of mitosis to promote centriole disengagement (Matsuo et al., 2011; Lee and Rhee, 2012; Kim et al., 2015); it influences the DNA Damage Response (DDR) (Wang et al., 2013; Antonczak et al., 2016).

Pericentrin defects have been linked to several human diseases (Delaval and Doxsey, 2010). Most importantly, mutations in human Pericentrin cause microcephalic osteodysplastic primordial dwarfism (MOPD) or Seckel syndrome, diseases associated with severe growth retardation during both foetal and post-foetal development (Rauch et al., 2008; Griffith et al., 2008; Bober and Jackson, 2017). Moreover, Pericentrin dysfunction has been linked to several other diseases such as mental disorders (Anitha et al., 2009), and diabetes (Huang-Doran et al., 2011). In none of these cases, however, is it understood how Pericentrin defects contribute to these complex pathologies.

Clearly it is important to understand Pericentrin function within the context of a developing organism, but such an analysis is complicated in vertebrates because centrosome and cilia defects lead to pleiotropic organismal defects. In vertebrates, the loss of centrosomes activates a p53-dependent pathway leading to cell death or senescence (Lambrus and Holland, 2017), while the loss of cilia leads to severe developmental defects (Mitchison and Valente, 2017). *Drosophila* is an attractive model system for this type of study, as flies can proceed quite normally through most of development in the absence of centrioles, centrosomes and cilia (Basto et al., 2006).

Here we examine *Drosophila* PLP function in the Sensory Organ Precursor (SOP) lineage of the pupal notum (Hartenstein and Posakony, 1989), and in testes —tissues in which the centrioles initially form centrosomes and then form cilia (Lattao et al., 2017). Using a combination of live cell imaging, Electron Microscopy (EM) and Electron Tomography (ET) we show that PLP helps to stabilise centrioles and to nucleate centriolar interphase MTs, and it promotes centriole cohesion, and centriole docking at the plasma membrane.

We also show that mother centrioles organise electron-dense “ pericentriolar clouds” that extend outwards from the cartwheel spokes and surround the centriolar MT triplets. These clouds are greatly diminished in *PLP* mutant centrioles, and we propose that the partial loss of these structures may explain the pleiotropic centriole, centrosome and cilium defects observed in *PLP* mutant cells.

## Results

### Centrioles separate prematurely in *PLP* mutant SOPs

To gain a better understanding of the role of PLP in centrosome organisation during mitosis we imaged the first cell division of either *WT* (Figure 1A) or strongly-hypomorphic/null *PLP* mutant (Figure 1B) SOPs. *PLP* mutant SOPs fell into two categories:(1) 10/17 SOPs (∼ 59%) entered mitosis with two centrosomes but, in 5/10 of these cells, centriole separation occurred prematurely compared to WT (arrows, Figure 1A, t=15 and Figure 1B, t=9; Figure 1F); (2) 7/1 7 SOPs (41%) entered mitosis with centrioles that had already prematurely separated (arrow, Figure 1C, t=-36; Figure 1F). These cells formed multipolar spindles as they entered mitosis (*yellow* arrow, Figure C, t=6),1 but became bipolar before anaphase onset (Figure 1C, t=12). Many of the centrosomes that had separated prematurely appeared to duplicate (*red* arrows, Figure 1C, t=12), so daughter cells often had too many centrosomes (*red* arrow, Figure 1, t=36; Figure 1G). Surprisingly, *PLP* mutant SOPs were not delayed in mitosis (Figure 1D,E), and there were no detectable defects in spindle alignment relative to the anterior-posterior body axis (Figure 1H). Thus, cell division is relatively unperturbed in *PLP* mutant SOPs, but centrioles often separate prematurely, leading to spindle multipolarity and centriole amplification.

**Figure 1:**
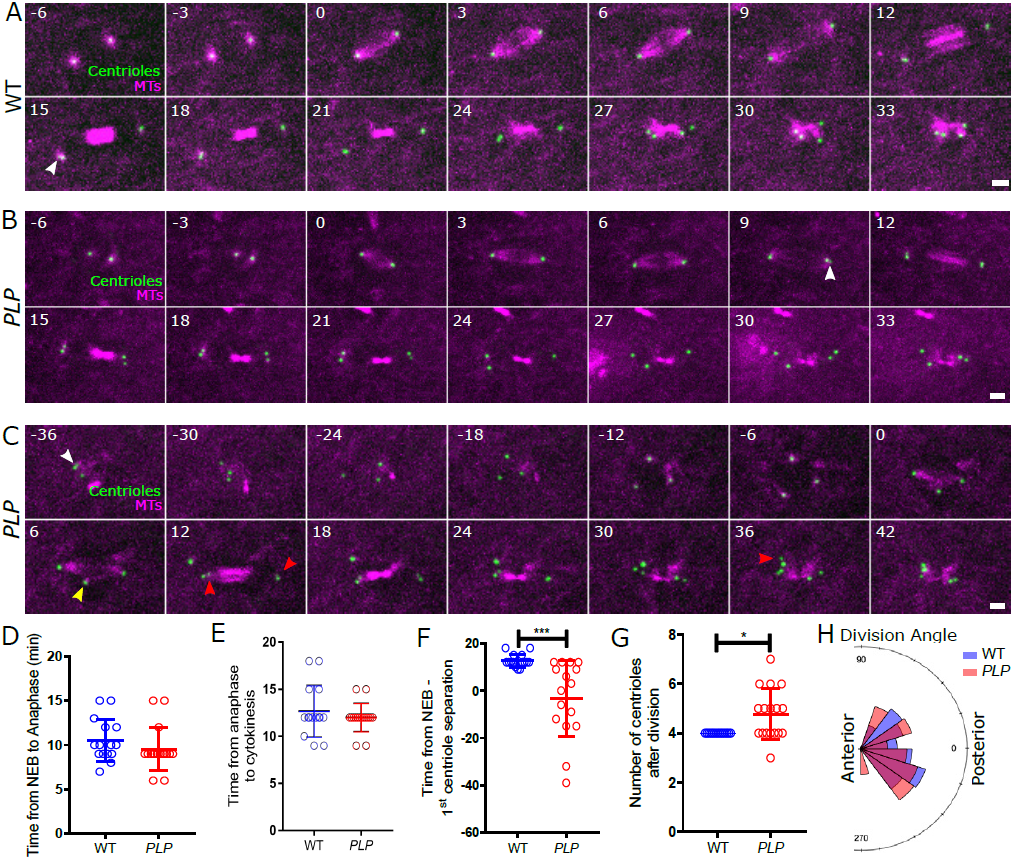
Mitosis is not dramatically perturbed in *PLP* mutant SOPs, but centrioles can separate prematurely. (**A-C**) Images from videos of living WT (A) or *PLP* mutant (B,C) SOPs expressing Jupiter-mCherry to reveal the MTs (*magenta*) and Asl to reveal the centrioles (*green*). Time in minutes relative to nuclear envelope breakdown (NEB) (t=0) is indicated; scale bar = 2μm. *White* arrowheads indicate when centriole separation is first detected; *yellow* arrowheads indicate an extra spindle pole; *red* arrowheads indicate instances where the centrioles in cells with extra centrioles separate. (**D-G**) Graphs compare various aspects of the behaviour of WT (*blue*) and *PLP* mutant (*red*) cells, as indicated. (**H**) Chart shows the division angle relative to the anterior/posterior axis of WT and PLP mutant SOPs. Distributions were assessed by the D’Agostino & Pearson normality test. Significance test for normal distributions was made by an unpaired two tailed T-Students test and for non-normal distribution by Mann-Whitney ranking test. All significance tests shown in this and subsequent Figures were performed in this manner unless specified. * p < 0.05, *** p < 0.001. Information on numbers analysed and biological repeats for these and all other experiments is given in Table S1. In (G) a Wilcoxon signed-rank test was used to compare the median of *PLP* mutant to the WT value of 4.

### The apical positioning of centrioles is perturbed in *PLP* mutant SOPs

When viewed along the apical-basal axis, the spindles in WT SOPs were well aligned with the cortex during metaphase—early-anaphase (Figure 2A, t=6- 12), but the posterior centriole pair initially moved basally during late-anaphase—telophase (t=15-8mins) -, before moving back to the apical cortex as the centrioles separated (t=21-27)—as described previously (Jauffred et al., 2013). At the end of mitosis all 4 centrioles were tightly clustered at the cortex, close to the spindle mid-body-remnant (t=30-33). In *PLP* mutant SOPs the spindles aligned with the cortex during metaphase—early anaphase (Figure 2B, t=61), and the posterior centriole pair moved basally during late-anaphase—telophase (t=15-18min), but the movement of the centrioles back to the apical cortex was often delayed and erratic (Figure 2B, t=21-33; Figure 2C,D).

**Figure 2:**
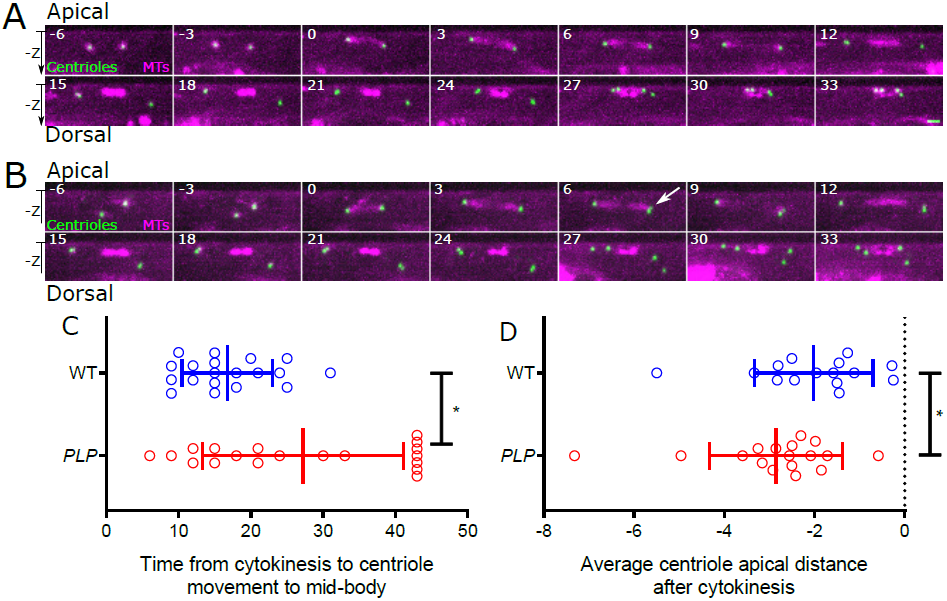
Centrioles do not migrate properly to the cortex at the end of cell division in *PLP* mutant SOPs. (**A,B**) Images from videos of living WT (A) or *PLP* mutant (B) SOPs expressing Jupiter-mCherry to reveal the MTs (*magenta*) and Asl to reveal the centrioles (*green*). Time in minutes relative to nuclear envelope breakdown (NEB) (t=0) is indicated; scale bar = 2mm. These images are taken from the same videos shown in Figure 1A,B, but shown from a side-on view to the spindle. *White* arrow indicates a centriole pair separating prematurely. (**C,D**) Graphs compare the behaviour of WT (*blue*) and *PLP* mutant (*red*) cells, as indicated. * p < 0.05.

### PLP helps to organise electron-dense “ prericentriolar cloudsȍ that surround the mother centriole

These live-cell studies suggest that *PLP* mutant SOP centrioles exhibit two independent defects: (1) Mother and daughter centrioles separate prematurely either prior to mitosis (potentially allowing centrioles to overduplicate), or during mitosis; (2) The centrioles do not migrate normally to the apical cortex at the end of mitosis. To better understand the nature of these defects we analysed centriole ultrastructure by electron tomography (ET) in the pupal-notum (at 24hr after pupae formation [APF], when all mitotic divisions in the SOP lineage are completed) and also in 3^rd^ instar larval wing-discs—a tissue where centriole ultrastructure is better preserved than in the pupal-notum (unpublished observations). We analysed 8 WT centrioles (all single centrioles) and 18 *PLP* mutant centrioles (5 single, 4 pairs, and 9 aggregated centrioles) from the pupal notum, and 7 WT (6 pairs, 1single) and 5 *PLP* mutant (3 pairs, 2 single) centrioles from wing discs.

A striking feature of the tomograms of WT wing disc centrioles was the presence of “ cloudsȍ o1f electron dense material that surround the mother centriole; these pericentriolar clouds originated close to the outer spokes of the central cartwheel and extended outwards through the gaps between the MT blades (Figure 3A). Similar structures were present in WT notum centrioles, but structural details were hard to discern (Figure 3B). In *PLP* mutant centrioles, similar electron-dense clouds emanated from the cartwheel spokes and spread past the MT blades, but the outer regions that normally surround the mother centriole were greatly diminished. (Figure 3C-E). Importantly, previous super-resolution light microscopy studies identified diffuse clouds of PLP-fibrils organised around the mother centriole in an approximately nine-fold symmetric manner (Mennella et al., 201 2). Taken together, these data suggest that PLP is a major component of the outer region of these pericentriolar-clouds.

**Figure 3:**
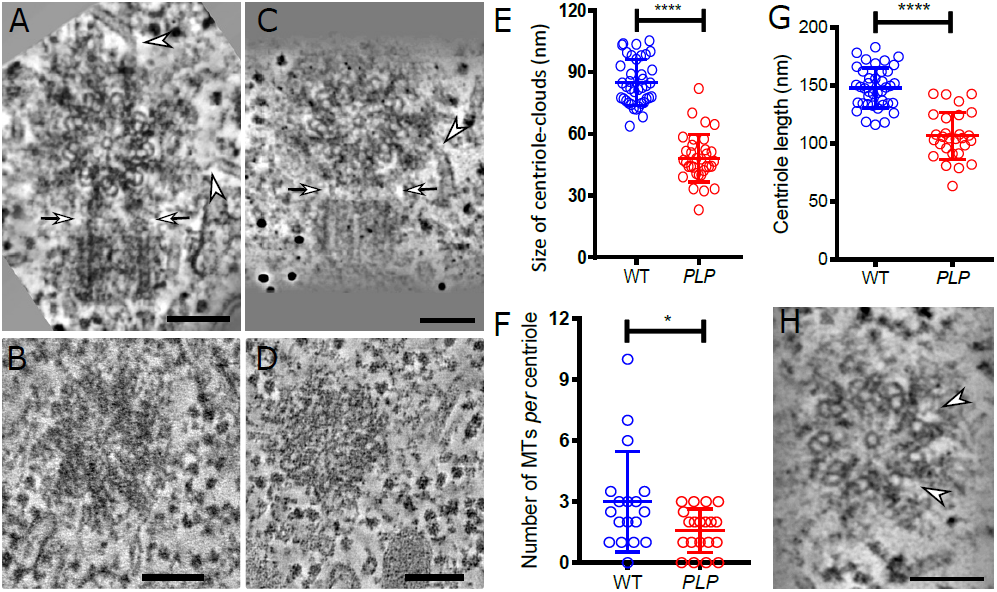
*PLP* mutants assemble reduced “ centriole-clouds” around the mother centriole. (**A-D**) Images from electron tomograms (ETs) of WT (A,B) or *PLP* mutant (C,D) centrioles in wing disc (A,C) or pupal notum (B,D) cells. Arrows in (A) highlight the electron-dense “ clouds” that spread outwards from the cartwheel spokes of the mother, through the gaps between the centriolar MTs, and contact the proximal end of the daughter centriole. The arrows in (C) highlight the same clouds that are present, but are greatly diminished in size, in *PLP* mutant centrioles. Arrowheads highlight cytoplasmic MTs that appear to be connected to the outer region of the centriole-clouds. (**E-G**) Graphs quantify either the distance that the centriole clouds spread outwards around the centrioles (E), the number of MTs associated with each centriole (F), or the length of the centrioles (G) in WT and *PLP* mutant cells. This data was combined from ET and EM analyses of both wing disc and pupal notum cells.(H) Electron tomogram image from a *PLP* mutant wingdisc centriole. Arrowheads highlight the missing outer B-MT. Scale bars = 100nm, * p < 0.05, **** p < 0.0001.

In WT centriole pairs, the pericentriolar centriole clouds directly contacted the proximal end of the daughter centrioles (n=6/6 centriole pairs examined) (arrows, Figure 3A). In *PLP* mutant centriole pairs (n=3/3), the daughter was still in contact with the mother, but the outer region of the fibrils was greatly reduced, so the daughter centriole was positioned much closer to the mother (arrows, Figure 3C). This raises the possibility that these pericentriolar clouds normally help promote mother/daughter centriole engagement, potentially explaining why *PLP* mutant centriole pairs tend to separate prematurely.

### PLP helps to organise centriolar MTs during interphase

Previous studies have shown that PLP promotes the organisation of the interphase PCM in cultured *Drosophila* cells (Mennella et al., 2012). Interestingly, MTs were often associated with the pericentriolar clouds (arrowheads, Figure 3A,C). To test if PLP was involved in organising interphase centriole MTs we counted the number of MTs within 100nm of the centriole MT wall (pooling data from both the pupal-notum and wing-disc). Significantly more MTs were associated with the centrioles in WT compared to *PLP* mutant tissues (Figure 3F). Thus, PLP helps organise interphase centriole MTs these cells, but *PLP* mutant centrioles can still associate with some MTs.

### Centrioles are too short and their organisation is subtly perturbed in the absence of PLP

Our EM analysis revealed two further aspects of the *PLP* mutant centriole phenotype. First, in two of the five singlet or mother centrioles we observed in PLP mutant wing discs several of the centriolar MT blades were missing an outer B-MT (arrows, Figure 3H)—something we have never observed in WT centrioles. This suggests that the PLP may help to assemble and/or stabilise the centriolar B-MTs. Second, centrioles were significantly shorter in the *PLP* mutant tissues (Figure 3G). Thus, centriole structure is slightly perturbed in *PLP* mutants.

### The inability of *PLP* mutant centrioles to interact with the cortex is correlated with their inability to nucleate interphase MTs

We measured the distance of the centrioles from the cell cortex in our ET images of the pupal notum and found that, in agreement with our live-cell studies, WT centrioles were closely associated with the cortex, while *PLP* mutant centrioles were more widely dispersed (Figure 4A-C). We counted the number of MTs associated with each of these centrioles and found that *PLP* mutant centrioles that associated with more MTs tended to be located closer to the cortex (Figure 4D). Thus, an inability to efficiently organise interphase MTs could contribute to the failure of mutant centrioles to become properly positioned at the cell cortex.

**Figure 4:**
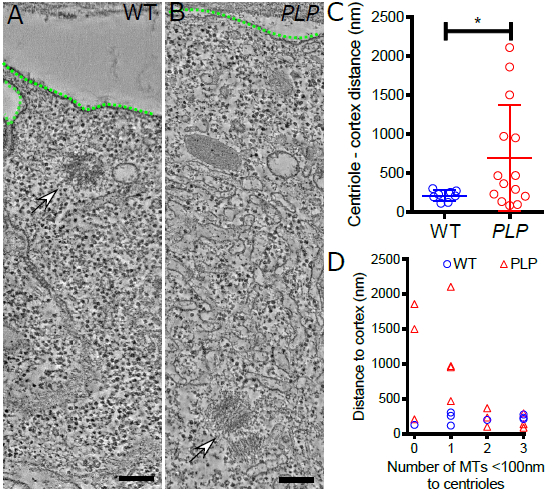
Centrioles are not closely associated with the cortex in *PLP* mutant cells. (**A,B**) Images from electron tomograms (ETs) of WT (A) or *PLP* mutant (B) pupal notum cells, highlighting the position of the centrioles (arrows) relative to the cell cortex (dotted *green* line). Scale bars = 100nm. (**C,D**) Graphs quantify either the centriole-to-cortex distance (C), or plot the number of MTs associated with each centriole relative to the distance of each centriole from the cortex (D). This data was analysed only in pupal notum cells. * p < 0.05.

### PLP mutant sensory cells lack cilia due to a failure to establish and/or maintain the position of the centrioles at the cell cortex

A lack of PLP/Pericentrin leads to defects in cilia function in flies and vertebrates (Martinez-Campos et al., 2004; Jurczyk et al., 2004). We therefore used Serial Block Face-Scanning Electron Microscopy (SBF-SEM) to reconstruct 3 notum bristle sensory organs from 72hr APF WT and *PLP* mutants (a time when organ assembly is complete). These organs had a complex cellular organisation, but this was not detectably perturbed in *PLP* mutants (Figure 5A,B). However, while centrioles, a transition zone (TZ) and a ciliary axoneme were all detectable in the 3 WT organs, none of these structures were detectable in the 3 *PLP* mutant organs (Figure S1). This suggests that cilia fail to form in *PLP* mutants because the centrioles are not correctly positioned within the sensory neurons.

**Figure 5:**
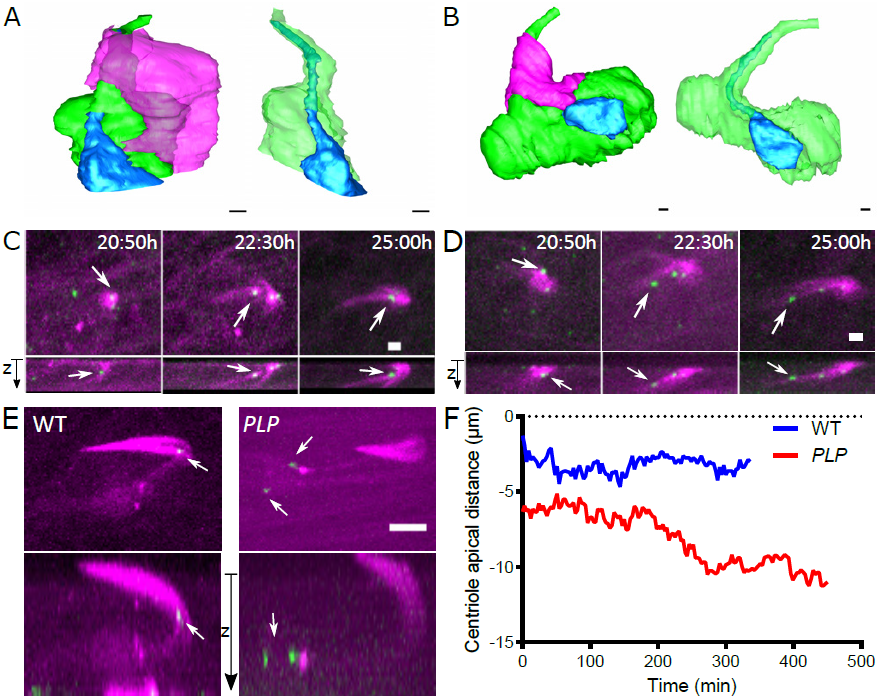
Basal bodies appear to be specified properly in *PLP* mutant sensory neurons, but they are mis-positioned. (**A,B**) Images show 3D- reconstructions from SBF-SEM data of the cells in a WT (A) or *PLP* mutant (B) pupal notum sensory organ. The Sensory Neuron (*blue*) sends an extension (that would normally contain the axoneme close to its tip) through the cell body of the Bristle Cell (*green*); the Support Cell (*magenta*) is illustrated in the images, on the left, but not on the right, which are also rotated by 90^o^. The overall organisation of the *PLP* mutant organ is not detectably perturbed. (**C-E**) Images from videos of living WT (C, E) or *PLP* mutant (D - E) sensory organs expressing Jupiter-mCherry to reveal the MTs (*magenta*) and Cep-104-GFP to reveal the centrioles and basal bodies (*green*). Time in hours:minutes after puparium formation (APF) (t=0) is indicated; scale bar = 500nm (A, B) or 2μm (C,D) or 5μm (E). (**F**) Graph charts the position of the brightest Cep-104-GFP containing centriole (that will normally become the basal body) relative to the cortex in WT and *PLP* mutant sensory organs.

To better understand the reason for this centriole mis-positioning, we analysed centriole behaviour in the developing pupal-notum, starting at 20:50h APF when cell divisions in the SOP lineage are complete, but before the sensory cilia are starting to form. We labelled centrioles with Cep104- GFP, a protein that localizes to both centrioles and cilia (Satish Tammana et al., 2013; Jiang et al., 2012) (unpublished observations). In WT organs a prominent array of MTs starts to form at 20:50hr APF that is associated with the forming bristle cell; several centrioles are detectable at this stage, but the most brightly labelled centriole (that will form the basal body of the cilium) is slightly displaced below the MT array (arrow, Figure 5C, 20:50hr APF). By 22:30hr APF, this centriole has established a sub-apical position close to the bristle cell MT array, and this position is maintained as ciliogenesis initiates (arrow Figure 5C, 22:30-25:00hr APF; Figure 5E). In *PLP* mutant sensory organs this MT array is formed, and a prominently labelled centriole is initially detectable close to it (arrow, Figure 5D 20:50hr APF), but this centriole fails to establish a stable sub-apical position relative to the MT array, and it usually became further displaced as development proceeded (arrow, Figure 5D, 22:30-25:00hr APF; Figure 5E). At 30hr APF a single brightly labelled centriole pair remained positioned close to the MT array in WT sensory organs, but in *PLP* mutant organs this centriole pair was invariably displaced away from the MT array by several microns, and often it had prematurely separated (Figure 5E, F). Thus, a centriole pair destined to form the basal body is specified in *PLP* mutant sensory neurons, but it fails to establish and/or maintain its proper position within the neuron, and so ultimately cannot organise a cilium.

### *[H2]PLP* mutant spermatocyte centrioles are mis-oriented and fail to dock properly at the PM, but they form an axoneme and recruit TZ proteins

Apart from ciliated sensory neurons, the only other cell type that forms cilia/flagella in *Drosophila* are the cells of the sperm lineage (Lattao et al., 2017). The centrioles in primary spermatocytes are longer than in other fly tissues (∼ 1μ m compared to ∼ 100-150nm) and they grow short cilia from both mother and daughter centrioles (Gonzalez et al., 1998; Lattao et al., 2017). After meiosis, these centrioles will form the basal bodies of the sperm flagellum. WT spermatocytes contain 2 centriole pairs, but there are usually too many centrioles in *PLP* mutant spermatocytes and these are often short and fragmented (Figure 6A-C), as reported previously (Martinez-Campos et al., 2004; Galletta et al., 2014).

**Figure 6:**
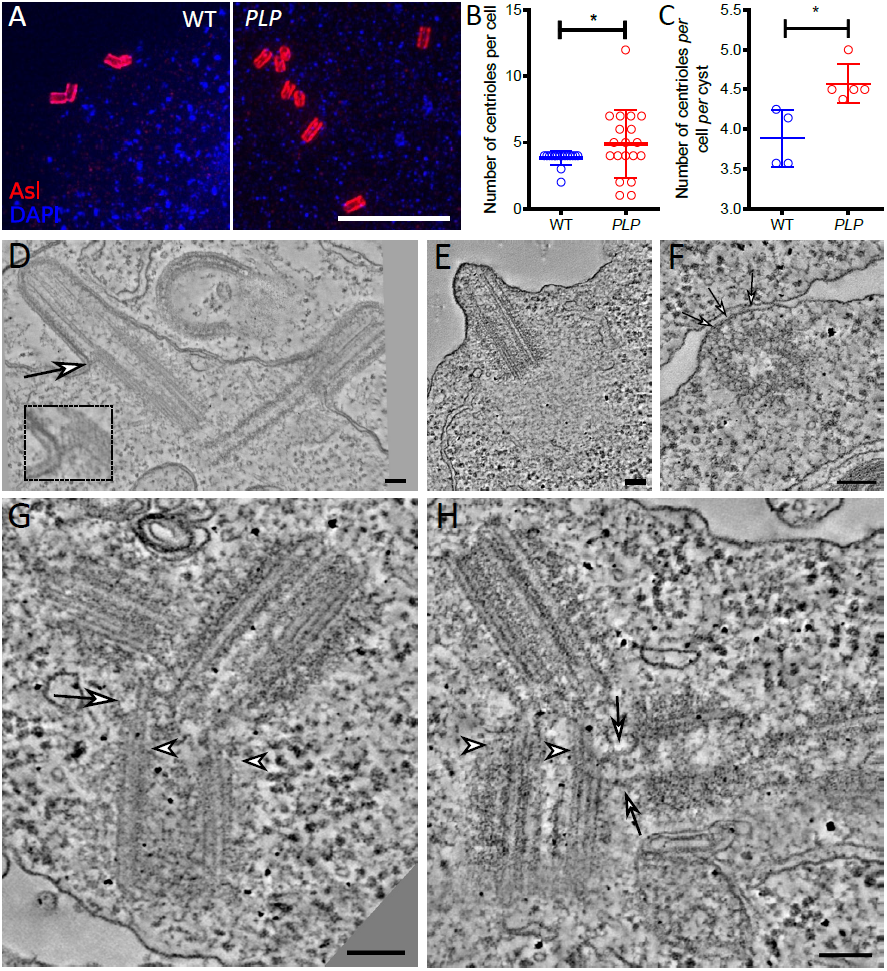
Centrioles and cilia show a range of defects in *PLP* mutant spermatocytes. (**A**) Images from fixed WT or *PLP* mutant spermatocytes stained with anti-Asl antibodies to reveal the centrioles (*red*) and DAPI to reveal the nuclei (*blue*). Scale bar = 5;Cm. (**B,C**) Graphs compare the number of centrioles per cell, or per cell per cyst (a cyst contains the 16 primary spermatocytes derived from the 4 rounds of division of the original spermatocyte gonialblast) in WT or *PLP* mutant testes. * p < 0.05. This analysis indicates that centrioles tend to overduplicate (as the number of centrioles per cell per cyst increases) but that centrioles can also mis- segregate, leading to individual cells with either too many or too few centrioles. (**D-H**) Micrographs show images from electron tomograms (ETs) of WT (D) or *PLP* mutant (E-H) spermatocytes. Scale bars = 100nm. Inset and arrows in (D) highlight an electron-dense region that we often observed connecting the centriole/basal body to the plasma membrane at the base of the cilium. Arrows in (F) highlight electron-dense regions that seem to connect the side of this mis-oriented centriole to the plasma membrane (PM). Arrows in G and H highlight regions where the clustered centrioles appear to connect to each other; arrowheads highlight that these centrioles appear to be forming an axoneme-like structure, where the centriole triplet MTs transition to axoneme-like doublet MTs.

An ET analysis revealed that WT centrioles (n=8) always organised a cilium, and the distal end of the centriole was often connected to the plasma membrane (PM), close to the position where the centriole MT triplets became axoneme MT doublets (inset, Figure 6D). In *PLP* mutant spermatocytes we found only a single centriole (113) that organised a cilium, and both the centriole and cilium were shorter than normal (Figure 6E). We also observed a single centriole (113) making a side-on, rather than end-on, connection to the plasma membrane (Figure 6F). In most cases, however, *PLP* mutant centrioles were found close to the PM, but were not detectably connected to the PM (Figure 6G,H). These centrioles were often clustered, and they usually formed an axoneme-like structure, as the MTs transitioned from triplets to doublets at the presumed distal ends (arrows, Figure 6G,H). Thus, *PLP* mutant centrioles can form short axoneme-like structures, but they very rarely form a cilium.

We next tested whether these axoneme-like structures can recruit TZ proteins. Surprisingly, all of the TZ proteins we examined were strongly recruited to the presumed distal ends of the *PLP* mutant centrioles in a manner very similar to that observed in WT centrioles that formed cilia (Figure 7A,B) (Vieillard et al., 2016; Pratt et al., 2016). Thus, *PLP* mutant centrioles appear to efficiently assemble axoneme- and TZ-like structures even though they are often not directly associated with the PM.

**Figure 7:**
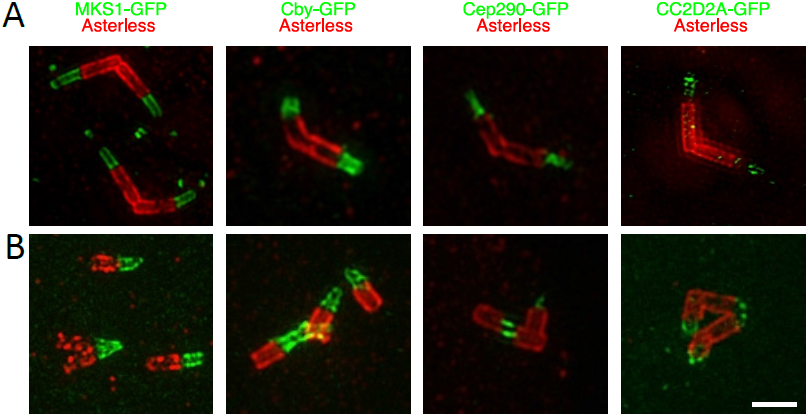
*PLP* mutant centrioles/basal bodies can recruit TZ proteins. (**A,B**) Micrographs show images from fixed WT (A) or *PLP* mutant (B) spermatocytes stained with anti-Asl antibodies to reveal the centrioles (*red*), and anti-GFP antibodies (*green*) to reveal the distribution of GFP-fusions to the TZ proteins MKS1, Cby, Cep290 and CC2D2A (as indicated). Scale bar = 1μm. Although EM studies show that the vast majority of *PLP* mutant centrioles are not connected to the PM, almost all of them appear to organise TZ proteins in a manner very similar to WT centrioles that are forming a cilium.

### PLP helps to organise interphase centriole MTs in spermatocytes

Our studies in the wing disc and notum indicate that PLP helps to organise interphase centriolar MTs, and this defect may contribute to the failure to position mutant centrioles at the apical surface. We therefore counted centriolar MT number in WT and *PLP* mutant spermatocytes by EM. We typically observed many MTs around WT centrioles (n=8), and these usually had their minus ends capped close to the centriole surface (Figure 8A-C). In *PLP* mutants (n=13) the number of MTs associated with the centrioles was greatly reduced, and we only observed a single MT that had its minus end capped at the centriole surface (Figure 8A; arrow, Figure 8D).

**Figure 8:**
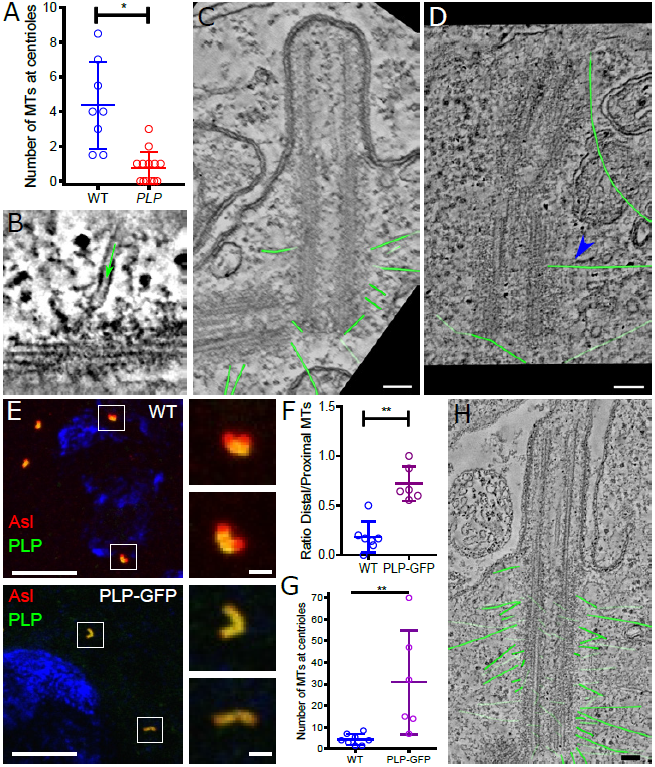
*PLP* helps to recruit MTs to the interphase centriole in spermatocytes. (**A**) Graph quantifies the number of cytoplasmic MTs associated with the centrioles in ETs of WT or PLP mutant spermatocytes. (**B-D**) Images from ETs of WT (B,C) or *PLP* mutant (D) spermatocytes: (*green* arrow, B) shows a MT with a “ capped” minus end attached to an electron-dense region on the outer wall of a WT centriole; (C,D) shows traces of all the cytoplasmic MTs (highlighted in *green*) associated with WT or PLP mutant centrioles. Only one MT is closely associated with the PLP mutant centriole (*blue* arrow, D). Scale bars = 100nm. (**E**) Images show fixed spermatocytes stained with anti-Asl antibodies to reveal the centrioles (*red*) and anti-PLP antibodies (*green*). The DNA in is stained with DAPI (*blue*), and the insets illustrate how PLP-GFP is distributed along the entire length of the centriole. Scale bars = 10mm (1mm in insets). **(F-H)** Graphs quantifying the MT ratio of MTs emanating from the distal versus proximal end of WT and PLP-GFP centrioles (F) and the MT numbers (G) and an image showing the centriole and traced MTs (*green*) (H). Scale bar = 100nm.

The endogenous PLP protein is normally concentrated proximally at spermatocyte centrioles (Galletta et al., 2014) (Figure 8E), and centriolar MTs were preferentially associated with the proximal half of these centrioles (Figure 8C and 8F). A GFP-PLP fusion protein expressed from the Ubq- promoter localises along the length of the spermatocyte centrioles (Figure 8G), and this led to an increase in the number of MTs associated with the centrioles (Figure 8G-H), and more of these MTs were associated with the centriole distal end (Figure 8F, H). These data strongly suggest that PLP is involved in organising interphase centriolar MTs in spermatocytes.

### The PCM proteins Spd-2 and Cnn help to recruit and/or maintain PLP at interphase centrioles, but antagonise the ability of PLP to organise MTs

In cultured cells PLP recruits PCM proteins such as Cnn and γ -tubulin to interphase centrioles (Mennella et al., 2012). *In vivo*, however, Cnn normally cooperates with Spd-2 to form a scaffold that recruits the mitotic PCM, and the role of these proteins in interphase PCM assembly is unclear (Conduit et al., 2015). We wondered, therefore, whether PLP functions to promote MT nucleation at interphase centrioles by recruiting Spd-2 and Cnn. Surprisingly, the centriolar levels of the PCM components Asl, Cnn, Spd-2, γ -tubulin and Polo-GFP were not detectably perturbed in *PLP* mutants (Figure 9A; Figure S2), indicating that PLP is not required to recruit and/or maintain these proteins at interphase spermatocyte centrioles.

**Figure 9:**
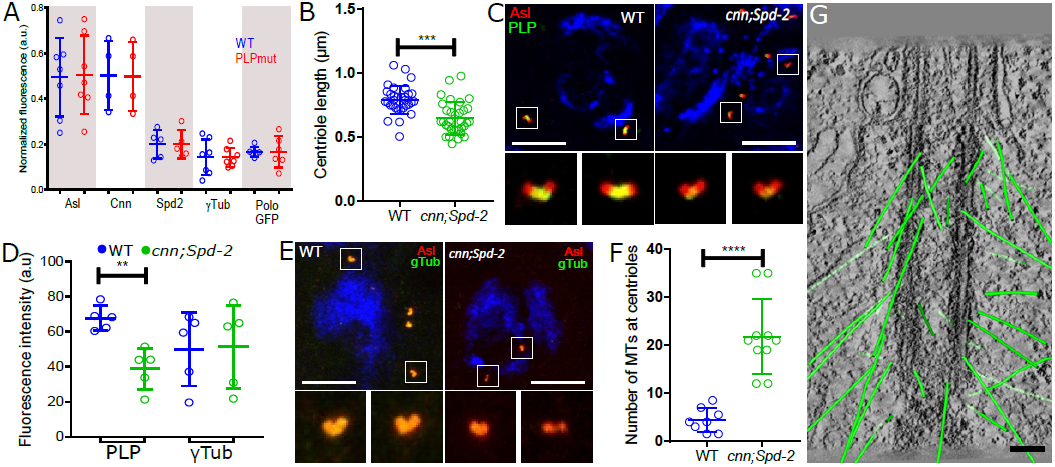
PLP does not recruit several PCM proteins to interphase spermatocyte centrioles, and Spd-2 and Cnn antagonise the ability of PLP to organise centriolar MTs. (**A**) Graph (images in Figure S2) quantifies the amount of various PCM proteins recruited to interphase centrioles in WT and *PLP* mutant spermatocytes. (B) Graph quantifies the length of interphase centrioles in *cnn*;*Spd-2* double mutant spermatocytes. (**C-E**) Images show (C,E) and graph quantifies (D) the amount of PLP or μ-tubulin (*green*) recruited to interphase centrioles in WT and *cnn*;*Spd-2* double mutant (right) spermatocytes (note that graphs show the levels of these proteins normalised to the amount of the centriole marker Asl, to allow for the centrioles being shorter in the mutant spermatocytes). (**F,G**) Graph quantifies (F) and image illustrates (G) the number of MTs associated with the centrioles in *cnn*;*Spd-2* double mutant spermatocytes. In (G) the MTs are highlighted (*green*)

Although PLP does not seem to recruit Spd-2 or Cnn to interphase spermatocyte centrioles, we reasoned that PLP might still promote interphase centriolar MTs by influencing their behaviour in some way. We therefore quantified PLP, g-Tub and MT nucleation in *cnn*;*Spd-2* double mutant spermatocytes (Figure 9B,C). Surprisingly, the centriolar levels of PLP, but not γ tubulin, were significantly reduced in the absence of Spd-2 and Cnn (Figure 9C-E), indicating that Spd-2 and Cnn play some part in recruiting and/or maintaining PLP at interphase centrioles. Even more surprisingly, the number of interphase centriolar MTs dramatically increased in the absence of Spd-2 and Cnn (Figure 9F,G). Thus, at least in spermatocytes, Spd-2 and Cnn do not promote MT organisation at interphase centrosomes, but rather they appear to suppress it.

## Discussion

Here we have used live-cell imaging combined with EM and ET to systematically catalogue the centriole, centrosome and cilium defects in *PLP* mutant pupal notum sensory organs and spermatocytes. We show that WT mother centrioles are normally surrounded by electron-dense clouds of material that extend outwards from the cartwheel spokes. In the absence of PLP, the outer region of these pericentriolar clouds is greatly reduced; this is in good agreement with previous studies that identified diffuse PLP-fibrils organised around the mother centriole in an approximately nine-fold symmetric manner (Mennella et al., 2012). We show that centrioles exhibit multiple, complex, centriole defects in the absence of PLP, several of which are in agreement with defects described in previous reports, but some of which have not been reported previously. Importantly, we propose that the loss of the outer-region of these pericentriolar clouds could explain most, if not all, of the defects we observe in *PLP* mutant centrioles.

The structure of centrioles is subtly perturbed in the absence of PLP: centrioles were too short and we occasionally observed centriole MT doublets that were missing an outer-B-MT. To our knowledge, these observations provide the first evidence that PLP is involved in maintaining centriole structure *per se*, and we suggest that the pericentriolar-clouds, which pass close by the centriolar MTs as they spread outwards from the central cartwheel, may have a role in helping to establish and/or maintain these B- MTs. It is unclear why centrioles are too short in the absence of PLP, but this could be a result of a general destabilisation of centriole structure. Mother and daughter centrioles tend to separate prematurely in the absence of PLP, and this can lead to centriole overduplication. Thus, PLP, like Pericentrin in vertebrates (Matsuo et al., 2012; Lee and Rhee, 2012; Kim et al., 2015), helps to maintain mother/daughter centriole cohesion. The outer regions of the pericentriolar clouds appear to connect mother and daughter centrioles, potentially explaining how PLP might contribute to cohesion. Importantly, this mechanism cannot be the only way that centriole cohesion is maintained in flies, as most centrioles still separated at the correct time in our live-cell analysis of SOPs, and we could still observe engaged mother and daughter centrioles in our EM and ET analysis, even when the pericentriolar clouds were greatly reduced. Thus, PLP contributes to mother/daughter centriole cohesion in flies, but is not essential for it. In vertebrates, PLP appears to be cleaved by Separase at the end of mitosis to promote centriole disengagement (Matsuo et al., 2012; Lee and Rhee, 2012); it will be fascinating to see if the outer-regions of the pericentriolar clouds are lost as centrioles disengage at the end of mitosis in fly cells.

It has previously been shown that PLP recruits PCM components to the interphase centriole in fly cultured cells (Mennella et al., 2012), but the role of PLP in organising interphase centriolar MTs has not been directly assessed. Our EM studies demonstrate that PLP plays an important part in organising these MTs; the number of interphase centriolar MTs decreased in the absence of PLP (although they were not abolished) and increased when PLP was overexpressed. Most surprisingly, however, several PCM proteins appear to be recruited to interphase spermatocyte centrioles normally in the absence of PLP, indicating that interphase centrioles likely use different mechanisms to recruit PCM in cultured cells and in spermatocytes. It is unclear how PLP promotes MT organisation at interphase centrioles, but our data strongly suggests that in spermatocytes it does not do so by recruiting or regulating Spd-2 and Cnn. These two proteins cooperate to form a scaffold that is essential for mitotic centrosome assembly in flies (Conduit et al., 2014; Feng et al., 2017), but they are also present at interphase centrioles (Mennella et al., 2012; Fu and Glover, 2012). Surprisingly, however, the ability of spermatocyte centrioles to organise MTs during interphase was dramatically increased in the absence of Spd-2 and Cnn. A possible explanation for this result is that Spd-2 and Cnn normally interact with PLP at interphase centrioles, but these interactions inhibit the ability of PLP to promote MT nucleation. Taken together, our findings suggest that interphase centrioles can organise MTs independently of key proteins that organise MTs at mitotic centrosomes, and that the organisation of interphase centriolar MTs may be complicated, and may use different mechanisms in different cell types.

In the absence of PLP, centrioles often fail to migrate properly to the cell cortex, and they fail to establish and/or maintain a proper connection to the PM in cells that form cilia. This presumably explains why PLP mutants exhibit severe cilia defects (Martinez-Campos et al., 2004). This is different to the situation in vertebrate cells where cilia lacking Pericentrin are also dysfunctional, but this has been attributed to a failure to recruit IFT and Polycystin2 proteins (Jurczyk et al., 2004). It is unclear why centrioles fail to migrate to, and/or establish a connection with, the PM in *PLP* mutant flies, but we noticed that mutant centrioles that associated with more MTs tended to be located closer to the apical cortex. Our sample size is small, so we remain cautious in interpreting this data, but it is consistent with the possibility that an inability to properly organise interphase MTs may contribute to the inability of the mutant centrioles to migrate properly to the apical cortex and to establish and/or maintain a connection with the PM. Another possibility is that the centriole clouds can directly or indirectly interact with PM and so help to establish and/or maintain the cortical position of the centrioles.

Finally, our analysis of centriole behaviour in *PLP* mutant spermatocytes revealed that even when these centrioles fail to connect properly to the PM, they can still assemble an axoneme-like structure. The centriolar MTs transitioned from a triplet to a doublet organisation at their presumed distal ends (as normally occurs when centrioles form an axoneme), and, most remarkably, the presumed distal regions recruited several TZ proteins that were organised in a very similar manner to WT TZs. The ability to recruit and organise TZ proteins is very surprising, as many of these proteins contain membrane-association domains, yet these centrioles are not associated with the PM. Thus, basal bodies appear capable of organising a TZ even when they fail to dock at the PM.

## Acknowledgements

We thank members of the Raff lab for valuable discussions. EM was performed at the Dunn School EM facility, part of the Micron Oxford Advanced Bioimaging Unit, funded by a Strategic Award from the Wellcome Trust (107457). The SBF-SEM system was funded by a BBSRC Alert 13 award to Chris Hawes (BB/L0141 22/1), M.B.P. was supported by a BBSRC PhD Studentship, H.R., and J.W.R. were supported by a Wellcome Trust Senior Investigator Award1 (104575).

## Materials and Methods

### Fly stocks

*w67* was used as a WT control in all experiments. Two previously described *PLP* mutant alleles were used in this study: plp ^272^ and plp^5^, both thought to be strong hypomorphic or null alleles (Martinez-Campos et al., 2004). Both alleles produced indistinguishable phenotypes and were used interchangeably. The *cnn;Spd-2* double mutant stock was created with the following lines: *cnn*^*hk21*^ (Megraw et al., 1999) and *cnn*^*f*04547^ (Lucas and Raff, 2007); *Spd-2*^*z*-5711^ and *Df(3L)st-j7* (Bloomington stock #5416) (Giansanti et al., 2008). UASt-GFP-Asterless (JWR and Renata Basto, unpublished) driven by the Scabrous–Gal4 transgene, a pan-neuronal Drosophila driver (Mlodzik et al., 1990) was used to mark the centrioles in the SOP lineage. The following transgenic lines were described previously: MKS1-GFP and CC2D2A-GFP (Pratt et al., 20^1^6), Cby-GFP (Enjolras et al., 2012), Cep290- GFP (Basiri et al., 2014), Polo-GFP (Buszczak et al., 2006) and PLP-GFP (Galletta et al., 2014).

### Antibodies

The following primary antibodies were used: Guinea-Pig anti-Asterless (Novak et al., 2014) (RaffLabDB#191); Rabbit anti-PLP (Martinez-Campos et al., 2004) (RaffLabDB#86), Rabbit anti-Cnn (Lucas and Raff, 2007) (RaffLabDB#37), Mouse anti-γTub (GTU88, Sigma, cat. T6557), Rabbit Anti-Spd-2 (Dix and Raff, 2007) (RaffLabDB#57). The following secondary antibodies were used: Anti-Guinea Pig IgG Alexa Fluor® 568 (Invitrogen, cat. A11075), Anti-Guinea Pig IgG Alexa Fluor® 633 (Invitrogen, cat. A21105),Anti-Rabbit IgG Alexa Fluor® 488 (Invitrogen, cat. A22106), Anti-Mouse IgG Alexa Fluor® 488 (Invitrogen, cat. A11001) and Anti-Mouse IgG Alexa Fluor® 568 (Invitrogen, cat. A11004).

### Fluorescence microscopy

3D-SIM of testis was performed as described previously (Roque et al., 2012). Live imaging of pupae centrioles and MTs was performed in a Nikon Eclipse TE200-E spinning disk confocal system, equipped with an EM-CCD Andor iXon+ camera, controlled by the Andor IQ2 software. Pupae were prepared for imaging as previously described (Jauffred et al., 2013). Testis squashes of whole testis and spermatocyte cysts were performed as previously reported (Roque et al., 2012; Dix and Raff, 2007). Immuno-fluorescence images were acquired in an Olympus Fluoview FV- 1000 microscope and software. Each slide imaged had both a control and a mutant testis in opposite sides of the slide to guarantee equal staining. Imaging conditions were maintained between slides.

### Image analysis

4D centriole tracking was performed in 8-bit converted images with Fiji (Schindelin et al., 2012) using the Trackmate plugin (Tinevez et al., 2016) using the following parameters: detection with sub-pixel localization using LOG method, threshold of 200 and estimated blob diameter set to 0.9;Cm. No initial threshold was applied to detections. The simple LAP tracker was used to create the tracks. Both linking max distance and gap-closing max distance were set to 2μm and Gap-closing max frame gap was set to 2. Centriole positions in 4D were extracted from Trackmate and exported into Prism (Graphpad) for analysis and plotting. Angles of division were calculated using the line function of Fiji and measured in relation to the anterior to posterior axis. Circular plots were computed using the circular package in Rstudio.Immuno-fluorescence images were analysed in Fiji, using a purposed-written macro (available on request) to automatically segment and extract the mean sum of marker fluorescence per centriole. Asterless staining was presented as mean sum per centriole, while all other markers where presented as ratio (marker mean sum)/(Asterless mean sum) per centriole. All data analysis and plotting was performed with Prism (Graphpad).

### Electron microscopy, tomography and Serial Block Face Scanning Electron Microscopy

Samples, processing and modelling of electron microscopy and tomography data of testis centrioles and pupal samples was performed as previously published (Roque et al., 2012; Pratt et al., 2016). Data was automatically acquired in an FEITecnai T12 at 120KV using a Gatan OneView digital camera with the Navigator function of SerialEM software (Kremer et al., 1996; Mastronarde and Held, 2017). For SBFSEM samples were prepared according to (Wilke et al., 2013) with some modifications. Briefly, samples were fixed overnight in 2.5% glutaraldehyde, 4% Paraformaldehyde and 0.1 % tannic acid at 4°;C (from a freshly prepared 10% stock) in 0.1 M PIPES buffer, pH 7.2. Samples were then washes twice for 30min in 0.1 M PIPES, followed by a 30 min wash in 50mM glycine in 0.1 M PIPES to quench free aldehydes, and another 30 min wash in 0.1 M PIPES. Samples were then embedded in 4% low melting point agarose plus 4% porcine gelatin (Melford, cat. L1204). Small cubes of agarose with one pupae each were then further fixed in1.5% potassium ferricyanide and 2% osmium tetroxide in 0.1 M PIPES for 1 hour at 4°;C. Samples were then washed three times for 10 min in water. Next samples were incubated in thiocarbohydrazide for 20min at room temperature, followed by three 10min washes in MQ water. Samples were then incubated in 2% osmium tetroxide in MQ water for 30min at 4C. After three 10 min MQ washes, samples were incubated in 1% uranyl acetate in MQ water overnight at 4°C. Samples were washed three times for 10 min in MQ water followed by *en-bloc* staining with lead aspartate solution for 30 min min at room temperature. After three 10 min washes in MQ water, dehydration was performed in ice with pre-cooled solutions of 30%, 50%, 70%, 90%, ^1^00%, 100% anhydrous ethanol for 10 min each, followed by ice-cold acetone for 10 min and a further 10 min in acetone at room temperature. Embedding was performed in acetone:Durcupan resin (Sigma cat. 44610) mix of 25% for 3 hours, 50% overnight, 75% for 3 hours and four times 100% Durcupan resin freshly prepared for 8-14 hours each. Samples were embedded in Beem capsules and cured at 60° C for 48 – 72 acquired using Digital Micrograph 2.0 every 50 nm, at 3.5KeV with 30 m aperture and VP set to 50 Pa. Images were aligned and modelled using the software package IMOD (Kremer et al., 1996). Iwata, K. Suzuki, Y. Sekine, H. Matsuzaki, M.

## Supplementary Figure Legends

**Figure S1.**
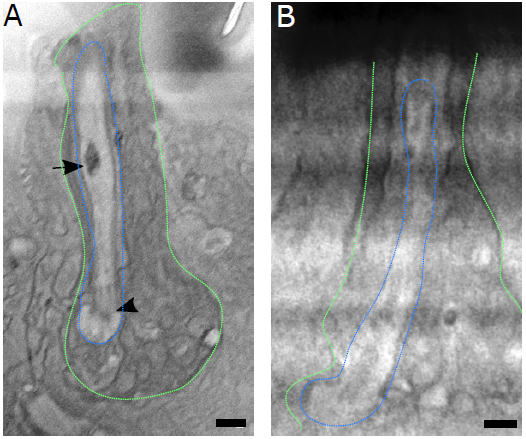
*PLP* mutant sensory cilia lack centrioles, TZs and axonemes. (A,B) Images from an SBF-SEM analysis of WT (A) or PLP mutant (B) Sensory Neurons (outlined by *green* dotted-line) showing the ciliary invagination (outlined by *blue* dotted-line). Although the resolution of these images is low the basal body can be observed at the base of the WT cilium (arrowhead), and the TZ can be seen as an electron-dense constriction of the PM just above the basal body. An electron-dense area of membrane that is a common feature of these type of cilia (ref) is highlighted (arrow). A ciliary invagination is present in the *PLP* mutant Sensory Neuron, but no centriole, TZ, axoneme or extra membrane are visible. Scale bar = 2μm.

**Figure S2.**
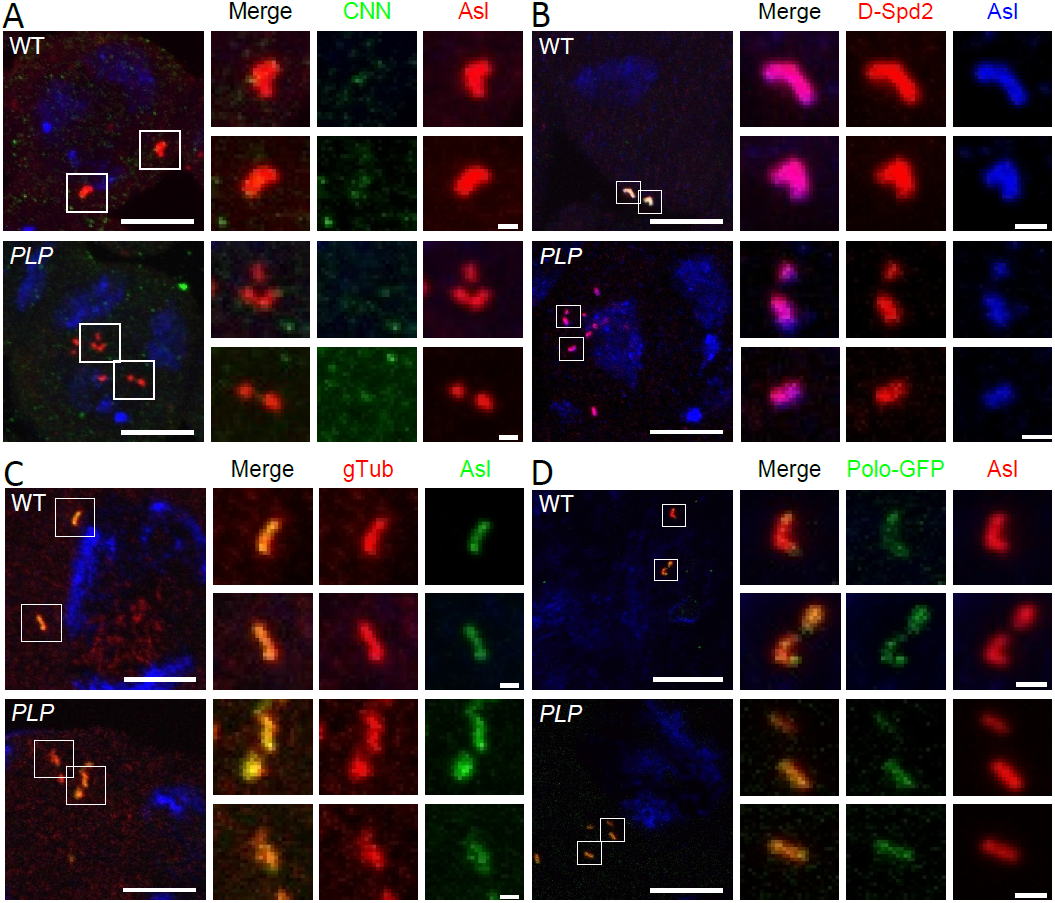
*PLP* mutant spermatocyte centrioles still recruit many PCM proteins. (**A-D**) Images show WT and *PLP* mutant (as indicated) spermatocyte centrioles stained with various centriole and PCM markers. Scale bar = 1μm, 1;Cm for insets.

## References

Alves-Cruzeiro, J.M.D.C., R. Nogales-Cadenas, and A.D. Pascual-Montano. 2013. CentrosomeDB: a new generation of the centrosomal proteins database for Human and Drosophila melanogaster. Nucleic Acids Research. 42:D430–6. doi:10.1093/nar/gkt1126.

Anitha, A., K. Nakamura, K. Yamada, Y. Iwayama, T. Toyota, N. Takei, Y. Iwata, K. Suzuki, Y. Sekine, H. Matsuzaki, M. Kawai, I. Thanseem, K. Miyoshi, T. Katayama, S. Matsuzaki, K. Baba, A. Honda, T. Hattori, S. Shimizu, N. Kumamoto, M. Kikuchi, M. Tohyama, T. Yoshikawa, and N. Mori. 2009. Association studies and gene expression analyses of theDISC1-interacting molecules, pericentrin 2 (PCNT2) and DISC1-binding zinc finger protein (DBZ), with schizophrenia and with bipolar disorder. Am. J. Med. Genet. B Neuropsychiatr. Genet. 150B:967–976.doi:10.1002/ajmg.b.30926.

Antonczak, A.K., L.I. Mullee, Y. Wang, D. Comartin, T. Inoue, L. Pelletier, and C.G. Morrison. 2016. Opposing effects of pericentrin and microcephalin onthe pericentriolar material regulate CHK1 activation inthe DNA damage response. Oncogene. 35:2003–2010. doi:10.1038/onc.2015.257.

Arquint, C., and E.A. Nigg. 2016. The PLK4-STIL-SAS-6 module at the core of centriole duplication. Biochem. Soc. Trans. 44:1253–1263. doi:10.1042/BST20160116.

Basiri, M.L., A. Ha, A. Chadha, N.M. Clark, A. Polyanovsky, B. Cook, and T. Avidor-Reiss. 2014. Ar ticleA Migrating Ciliary Gate Compartmentalizesthe Site of Axoneme Assembly in Drosophila Spermatids. Current Biology.24:1–10. doi:10.1016/j.cub.2014.09.047.

Basto, R., J. Lau, T. Vinogradova, A. Gardiol, C.G. Woods, A. Khodjakov, and J.W. Raff. 2006. Flies without centrioles. Cell. 125:1375–1386.doi:10.1016/j.cell.2006.05.025.

Bettencourt-Dias, M., F. Hildebrandt, D. Pellman, G. Woods, and S.A. Godinho. 2011. Centrosomes and cilia in human disease. Trends in Genetics. 27:307–315. doi:10.1016/j.tig.2011.05.004.

Bober, M.B., and A.P. Jackson. 2017. Microcephalic Osteodysplastic Primordial Dwarfism, Type II: a Clinical Review. Curr Osteoporos Rep.15:61–69. doi:10.1007/s11914-017-0348-1.

Buszczak, M., S. Paterno, D. Lighthouse, J. Bachman, J. Planck, S. Owen,A.D. Skora, T.G. Nystul, B. Ohlstein, A. Allen, J.E. Wilhelm, T.D. Murphy,R.W. Levis, E. Matunis, N. Srivali, R.A. Hoskins, and A.C. Spradling. 2006. The Carnegie Protein Trap Library: A Versatile Tool for Drosophila Developmental Studies. Genetics. 175:1505–1531. doi:10.1534/genetics.106.065961.

Conduit, P.T., A. Wainman, and J.W. Raff. 2015. Centrosome function and assembly in animal cells. Nat Rev Mol Cell Biol. 16:611–624.doi:10.1038/nrm4062.

Conduit, P.T., J.H. Richens, A. Wainman, J. Holder, C.C. Vicente, M.B. Pratt,C.I. Dix, Z.A. Novak, I.M. Dobbie, L. Schermelleh, and J.W. Raff. 2014. Amolecular mechanism of mitotic centrosome assembly in Drosophila. Elife.3:e03399. doi:10.7554/eLife.03399.

Delaval, B., and S.J. Doxsey. 2010. Pericentrin in cellular function and disease. J Cell Biol. 188:181–190. doi:10.1083/jcb.200908114.

Dix, C.I., and J.W. Raff. 2007. Drosophila Spd-2 recruits PCM to the sperm centriole, but is dispensable for centriole duplication. 17:1759–1764. doi:10.1016/j.cub.2007.08.065.

Enjolras, C., J. Thomas, B. Chhin, E. Cortier, J.-L. Duteyrat, F. Soulavie, M.J. Kernan, A. Laurençon, and B. Durand. 2012. Drosophila chibby is required for basal body formation and ciliogenesis but not for Wg signaling. J Cell Biol. 197:313–325. doi:10.1083/jcb.201109148.

Feng, Z., A. Caballe, A. Wainman, S. Johnson, A.F.M. Haensele, M.A. Cottee,P.T. Conduit, S.M. Lea, and J.W. Raff. 2017. Structural Basis for Mitotic Centrosome Assembly in Flies. Cell. 169:1078–1089.e13. doi:10.1016/j.cell.2017.05.030.

Fu, J., and D.M. Glover. 2012. Structured illumination of the interface between centriole and peri-centriolar material. Open Biol. 2:120104.doi:10.1098/rsob.120104.

Galletta, B.J., R.X. Guillen, C.J. Fagerstrom, C.W. Brownlee, D.A. Lerit, T.L. Megraw, G.C. Rogers, and N.M. Rusan. 2014. Drosophila pericentrin requires interaction with calmodulin for its function at centrosomes and neuronal basal bodies but not at sperm basal bodies. Mol Biol Cell. 25:2682–2694. doi:10.1091/mbc.E13-10-0617.

Giansanti, M.G., E. Bucciarelli, S. Bonaccorsi, and M. Gatti. 2008. Drosophila SPD-2 is an essential centriole component required for PCM recruitment and astral-microtubule nucleation. 18:303–309. doi:10.1016/j.cub.2008.01.058.

Gonzalez, C., G. Tavosanis, and C. Mollinari. 1998. Centrosomes and microtubule organisation during Drosophila development. J Cell Sci. 111 (Pt 18):2697–2706.

Griffith, E., S. Walker, C.-A. Martin, P. Vagnarelli, T. Stiff, B. Vernay, N. Al Sanna, A. Saggar, B. Hamel, W.C. Earnshaw, P.A. Jeggo, A.P. Jackson,and M. O'Driscoll. 2008. Mutations in pericentrin cause Seckel syndrome with defective ATR-dependent DNA damage signaling. Nat Genet.40:232–236. doi:10.1038/ng.2007.80.

Haren, L., T. Stearns, and J. Lüders. 2009. Plk1-dependent recruitment of gamma-tubulin complexes to mitotic centrosomes involves multiple PCM components. PLoS ONE. 4:e5976. doi:10.1371/journal.pone.0005976.

Hartenstein, V., and J.W. Posakony. 1989. Development of adult sensilla on the wing and notum of Drosophila melanogaster. Development. 107:389–405.

Huang-Doran, I., L.S. Bicknell, F.M. Finucane, N. Rocha, K.M. Porter, Y.C.L. Tung, F. Szekeres, A. Krook, J.J. Nolan, M. O'Driscoll, M. Bober, S. O'Rahilly, A.P. Jackson, R.K. Semple, Majewski Osteodysplastic Primordial Dwarfism Study Group. 2011. Genetic defects in human pericentrin are associated with severe insulin resistance and diabetes. Diabetes. 60:925–935. doi:10.2337/db10-1334.

Jauffred, B., F. Llense, B. Sommer, Z. Wang, C. Martin, and Y. Bellaiche. 2013. Regulation of centrosome movements by numb and the collapsing response mediator protein during Drosophila sensory progenitor asymmetric division. Development. 140:2657–2668. doi:10.1242/dev.087338.

Jiang, K., G. Toedt, S.M. Gouveia, N.E. Davey, S. Hua, B. van der Vaart, I. Grigoriev, J. Larsen, L.B. Pedersen, K. Bezstarosti, M. Lince-Faria, J. Demmers, M.O. Steinmetz, T.J. Gibson, and A. Akhmanova. 2012. A Proteome-wide Screenfor Mammalian SxIP Motif-Containing Microtubule Plus-End Tracking Proteins. Current Biology. 22:1800–1807. doi:10.1016/j.cub.2012.07.047.

Jurczyk, A., A. Gromley, S. Redick, J. San Agustin, G. Witman, G.J. Pazour,D.J.M. Peters, and S. Doxsey. 2004. Pericentrin forms a complex with intraflagellar transport proteins and polycystin-2 and is required for primary cilia assembly. J Cell Biol. 166:637–643. doi:10.1083/jcb.200405023.

Kim, J., K. Lee, and K. Rhee. 2015. PLK1 regulation of PCNT cleavage ensures fidelity of centriole separation during mitotic exit. Nat Commun. 6:10076. doi:10.1038/ncomms10076.

Kremer, J.R.J., D.N.D. Mastronarde, and J.R.J. McIntosh. 1996. Computer Visualization of Three-Dimensional Image Data Using IMOD. J Struct Biol. 116:6–6. doi:10.1006/jsbi.1996.0013.

Lambrus, B.G., and A.J. Holland. 2017. A New Mode of Mitotic Surveillance. Trends Cell Biol. 27:314–321. doi:10.1016/j.tcb.2017.01.004.

Lattao, R., L. Kovács, and D.M. Glover. 2017. The Centrioles, Centrosomes, Basal Bodies, and Cilia of Drosophila melanogaster. Genetics. 206:33–53.doi:10.1534/genetics.116.198168.

Lawo, S., M. Hasegan, G.D. Gupta, and L. Pelletier. 2012. Subdiffraction imaging of centrosomes reveals higher-order organizational features of pericentriolar material. Nat Cell Biol. 14:1148–1158. doi:10.1038/ncb2591.

Lee, K., and K. Rhee. 2011. PLK1 phosphorylation of pericentrin initiates centrosome maturation at the onset of mitosis. J Cell Biol. 195:1093–1101. doi:10.1083/jcb.201106093.

Lee, K., and K. Rhee. 2012. Separase-dependent cleavage of pericentrin B is necessary and sufficient for centriole disengagement during mitosis. Cell Cycle. 11:2476–2485. doi:10.4161/cc.20878.

Lerit, D.A., H.A. Jordan, J.S. Poulton, C.J. Fagerstrom, B.J. Galletta, M. Peifer, and N.M. Rusan. 2015. Interphase centrosome organization by the PLP-Cnn scaffold is required for centrosome function. J Cell Biol. 210:79– 97. doi:10.1083/jcb.201503117.

Lucas, E.P., and J.W. Raff. 2007. Maintaining the proper connection between the centrioles and the pericentriolar matrix requires Drosophila centrosomin. J Cell Biol. 178:725–732. doi:10.1083/jcb.200704081.

Martinez-Campos, M., R. Basto, J. Baker, M. Kernan, and J.W. Raff. 2004. The Drosophila pericentrin-like protein is essential for cilia/flagella function, but appears to be dispensable for mitosis. J Cell Biol. 165:673– 683. doi:10.1083/jcb.200402130.

Mastronarde, D.N., and S.R. Held. 2017. Automated tilt series alignment and tomographic reconstruction in IMOD. J Struct Biol. 197:102–113. doi:10.1016/j.jsb.2016.07.011.

Matsuo, K., K. Ohsumi, M. Iwabuchi, T. Kawamata, Y. Ono, and M. Takahashi. 2012. Kendrin Is a Novel Substrate for Separase Involved in the Licensing of Centriole Duplication. doi:10.1016/j.cub.2012.03.048.

Megraw, T.L., K. Li, L.R. Kao, and T.C. Kaufman. 1999. The centrosomin protein is required for centrosome assembly and function during cleavage in Drosophila. Development. 126:2829–2839.

Mennella, V., B. Keszthelyi, K.L. McDonald, B. Chhun, F. Kan, G.C. Rogers,B. Huang, and D.A. Agard. 2012. Subdiffraction-resolution fluorescence microscopy reveals a domain of the centrosome critical for pericentriolar material organization. Nat Cell Biol. 14:1159–1168. doi:10.1038/ncb2597.

Mitchison, H.M., and E.M. Valente. 2017. Motile and non-motile cilia in human pathology: from function to phenotypes. J. Pathol. 241:294–309. doi:10.1002/path.4843.

Mlodzik, M., N.E. Baker, and G.M. Rubin. 1990. Isolation and expression of scabrous, a gene regulating neurogenesis in Drosophila. Genes Dev.4:1848–1861. doi:10.1101/gad.4.11.1848.

Nigg, E.A., and J.W. Raff. 2009. Centrioles, centrosomes, and cilia in health and disease. Cell. 139:663–678. doi:10.1016/j.cell.2009.10.036.

Novak, Z.A., P.T. Conduit, A. Wainman, and J.W. Raff. 2014. Asterless licenses daughter centrioles to duplicate for the first time in Drosophila embryos. Curr. Biol. 24:1276–1282. doi:10.1016/j.cub.2014.04.023.

Palazzo, R.E., J.M. Vogel, B.J. Schnackenberg, D.R. Hull, and X. Wu. 2000.Centrosome maturation. Curr Top Dev Biol. 49:449–470.

Pratt, M.B., J.S. Titlow, I. Davis, A.R. Barker, H.R. Dawe, J.W. Raff, and H. Roque. 2016. Drosophila sensory cilia lacking MKS proteins exhibit striking defects in development but only subtle defects in adults. J Cell Sci. 129:3732–3743. doi:10.1242/jcs.194621.

Purohit, A., S.H. Tynan, R. Vallee, and S.J. Doxsey. 1999. Direct interaction of pericentrin with cytoplasmic dynein light intermediate chain contributes to mitotic spindle organization. J Cell Biol. 147:481–492.

Rauch, A., C.T. Thiel, D. Schindler, U. Wick, Y.J. Crow, A.B. Ekici, A.J. van Essen, T.O. Goecke, L. Al-Gazali, K.H. Chrzanowska, C. Zweier, H.G. Brunner, K. Becker, C.J. Curry, B. Dallapiccola, K. Devriendt, A. Dörfler, E. Kinning, A. Megarbane, P. Meinecke, R.K. Semple, S. Spranger, A. Toutain, R.C. Trembath, E. Voss, L. Wilson, R. Hennekam, F. de Zegher, H.-G. Dörr, and A. Reis. 2008. Mutations in the pericentrin (PCNT) gene cause primordial dwarfism. Science. 319:816–819. doi:10.1126/science.1151174.

Richens, J.H., T.P. Barros, E.P. Lucas, N. Peel, D.M.S. Pinto, A. Wainman, and J.W. Raff. 2015. The Drosophila Pericentrin-like-protein (PLP) cooperates with Cnn to maintain the integrity of the outer PCM. Biology Open. 4:bio.012914–1061. doi:10.1242/bio.012914.

Roque, H., A. Wainman, J. Richens, K. Kozyrska, A. Franz, and J.W. Raff. 2012. Drosophila Cep135/Bld10 maintains proper centriole structure but is dispensable for cartwheel formation. J Cell Sci. 125:5881–5886. doi:10.1242/jcs.113506.

Satish Tammana, T.V., D. Tammana, D.R. Diener, and J. Rosenbaum. 2013. Centrosomal protein CEP104 (Chlamydomonas FAP256) moves to the ciliary tip during ciliary assembly. J Cell Sci. 126:5018–5029. doi:10.1242/jcs.133439.

Schindelin, J., I. Arganda-Carreras, E. Frise, V. Kaynig, M. Longair, T. Pietzsch, S. Preibisch, C. Rueden, S. Saalfeld, B. Schmid, J.-Y. Tinevez, D.J. White, V. Hartenstein, K. Eliceiri, P. Tomancak, and A. Cardona. 2012. Fiji: an open-source platform for biological-image analysis. Nat Methods. 9:676–682. doi:10.1038/nmeth.2019.

Sonnen, K.F., L. Schermelleh, H. Leonhardt, and E.A. Nigg. 2012. 3Dstructured illumination microscopy provides novel insight into architecture of human centrosomes. Biology Open. 1:965–976.

Tinevez, J.-Y., N. Perry, J. Schindelin, G.M. Hoopes, G.D. Reynolds, E. Laplantine, S.Y. Bednarek, S.L. Shorte, and K.W. Eliceiri. 2016. TrackMate: An open and extensible platform for single-particle tracking.Methods. 115:80–90. doi:10.1016/j.ymeth.2016.09.016.

Vieillard, J., M. Paschaki, J.-L. Duteyrat, C. Augière, E. Cortier, J.-A. Lapart, J. Thomas, and B. Durand. 2016. Transition zone assembly and its contribution to axoneme formation in Drosophila male germ cells. J Cell Biol. 214:875–889. doi:10.1083/jcb.201603086.

Wang, Y., T.J. Dantas, P. Lalor, P. Dockery, and C.G. Morrison. 2013. Promoter hijack reveals pericentrin functions in mitosis and the DNA damage response. Cell Cycle. 12:635–646. doi:10.4161/cc.23516.

Wilke, S.A., J.K. Antonios, E.A. Bushong, A. Badkoobehi, E. Malek, M. Hwang, M. Terada, M.H. Ellisman, and A. Ghosh. 2013. Deconstructing complexity: serial block-face electron microscopic analysis of the hippocampal mossy fiber synapse. J. Neurosci. 33:507–522. doi:10.1523/JNEUROSCI.1600-12.2013.

Zimmerman, W.C., J. Sillibourne, J. Rosa, and S.J. Doxsey. 2004. Mitosis-specific anchoring of gamma tubulin complexes by pericentrin controls spindle organization and mitotic entry. Mol Biol Cell. 15:3642–3657. doi:10.1091/mbc.E03-11-0796.

